# Application of Optogenetic Amyloid-β Distinguishes Between Metabolic and Physical Damage in Neurodegeneration

**DOI:** 10.1101/836593

**Authors:** Lim Chu Hsien, Prameet Kaur, Emelyne Teo, Vanessa Lam, Fangchen Zhu, Caroline Kibat, Ajay Mathuru, Jan Gruber, Nicholas S. Tolwinski

## Abstract

The brains of Alzheimer’s Disease patients show a decrease in brain mass and a preponderance of extracellular Amyloid-β plaques. These plaques are formed by aggregation of polypeptides that are derived from Amyloid Precursor Protein (APP). Amyloid-β plaques are thought to play either a direct or an indirect role in disease progression, however the exact role of aggregation and plaque formation in the ethology of Alzheimer’s Disease is subject to debate, not least because the biological effects of soluble and aggregated Amyloid-β peptides are difficult to separate *in vivo*. To investigate the consequences of formation of Amyloid-β oligomers in living tissues, we developed a fluorescently tagged, optogenetic Amyloid-β peptide that oligomerizes rapidly in the presence of blue light. We applied this system to the crucial question of how intracellular Amyloid-β oligomers underlie the pathologies of Alzheimer’s Disease. We show that, although both expression and induced oligomerization of Amyloid-β were detrimental to lifespan and healthspan, we were able to separate the metabolic and physical damage caused by light-induced Amyloid-β oligomerization from Amyloid-β expression alone. The physical damage caused by Amyloid-β oligomers also recapitulated the catastrophic tissue loss that is a hallmark of late AD. We show that the lifespan deficit induced by Amyloid-β oligomers was reduced with Li^+^ treatment. Our results present the first model to separate different aspects of disease progression.

## Introduction

Alzheimer’s disease (AD) is a debilitating, age-associated, neurodegenerative disease which affects more than 46.8 million people worldwide and represents the 6th leading cause of death in the United States of America (Ahn et al. 2001, Zhang et al. 2011, De-Paula et al. 2012, Kumar et al. 2015, Hawkes 2016). Despite extensive efforts over the last 50 years, no disease-modifying therapy has been found and, most recently, several high-profile Phase III clinical trials have failed (Anderson et al. 2017, Lim et al. 2018, Park et al. 2018). AD clinical trial landscape has largely been dominated by the amyloid cascade hypothesis, with more than 50% of the drugs targeting amyloid beta (Aβ) in Phase III trial alone (Cummings et al. 2018). The amyloid cascade hypothesis, in its original form, posits the deposition of Aβ, in particular the extracellular Aβ plaque, as the main driver of AD (Hardy et al. 1992). However, the causative role of Aβ plaque has been challenged recently, not least because of the failure of numerous interventions targeting Aβ plaque in Phase III trials (Cummings 2018) and the observation of Aβ plaques in brains of non-AD symptomatic individuals. Model organisms ranging from nematodes to mice have further shown that AD pathology can be modeled in the absence of obvious Aβ plaques (Duff et al. 1996, Fong et al. 2016). These observations point to a different mechanism for Aβ neurotoxicity perhaps through Aβ’s intracellular accumulation (LaFerla et al. 2007).

Amyloid plaques are macroscopic extracellular protein aggregates, but intracellular Aβ must function in smaller units of single peptides, oligomers or aggregates. Of these, soluble Aβ oligomers appear to be the main toxic species in AD (Ferreira et al. 2011). For example, Aβ oligomers have been shown to induce neurotoxic effects including memory loss in transgenic animal models (reviewed in (Mroczko et al. 2018)). These studies, however, mostly rely on *in vitro* injection of synthetic Aβ oligomers or oligomers extracted from AD brains (Mroczko, Groblewska et al. 2018). There is currently a lack of tools that can directly control Aβ oligomerization *in vivo*, which will allow direct examination of the effects of Aβ oligomerization.

Here we describe an optogenetic method to study Amyloid-β (Aβ) protein oligomerization *in vivo* in different model organisms and apply it to delineating mechanisms of disease progression. Optogenetics hinges on light responsive proteins that can be fused to genes of interest, allowing for the spatial and temporal regulation of proteins in a highly precise manner. Temporal-spatial control is achieved simply by the exposure of the target system to lights of a specific wavelength, without the need to introduce other external agents (Fenno et al. 2011). One such optogenetic protein, a modified version of the *Arabidopsis* cryptochrome2 (CRY2) protein, oligomerizes quickly and reversibly in the presence of blue light at 488nm. CRY2 fusion proteins have, for instance, been used to study cortical actin dynamics in cell contractility during tissue morphogenesis, the dynamics of Wnt and EGF signaling, plasma membrane composition and transcriptional regulation (Kennedy et al. 2010, Idevall-Hagren et al. 2012, Guglielmi et al. 2015, Huang et al. 2017, Johnson et al. 2017, Kaur et al. 2017).

To address the role of aggregation of intracellular Aβ in the pathophysiological effects of AD, we developed a light-inducible system for use in model organisms. Specifically, we generated an *in vivo*, light-dependent, oligomerization switch for the formation and dissolution of Aβ oligomers in *Drosophila melanogaster, Caenorhabditis elegans* and *Danio rerio* (Figure 1a). This optogenetically-driven model allowed the visualization of microscopic Aβ oligomerization and addressed the question of their relationship to Aβ pathogenesis with spatiotemporal precision.

**Figure 1.**
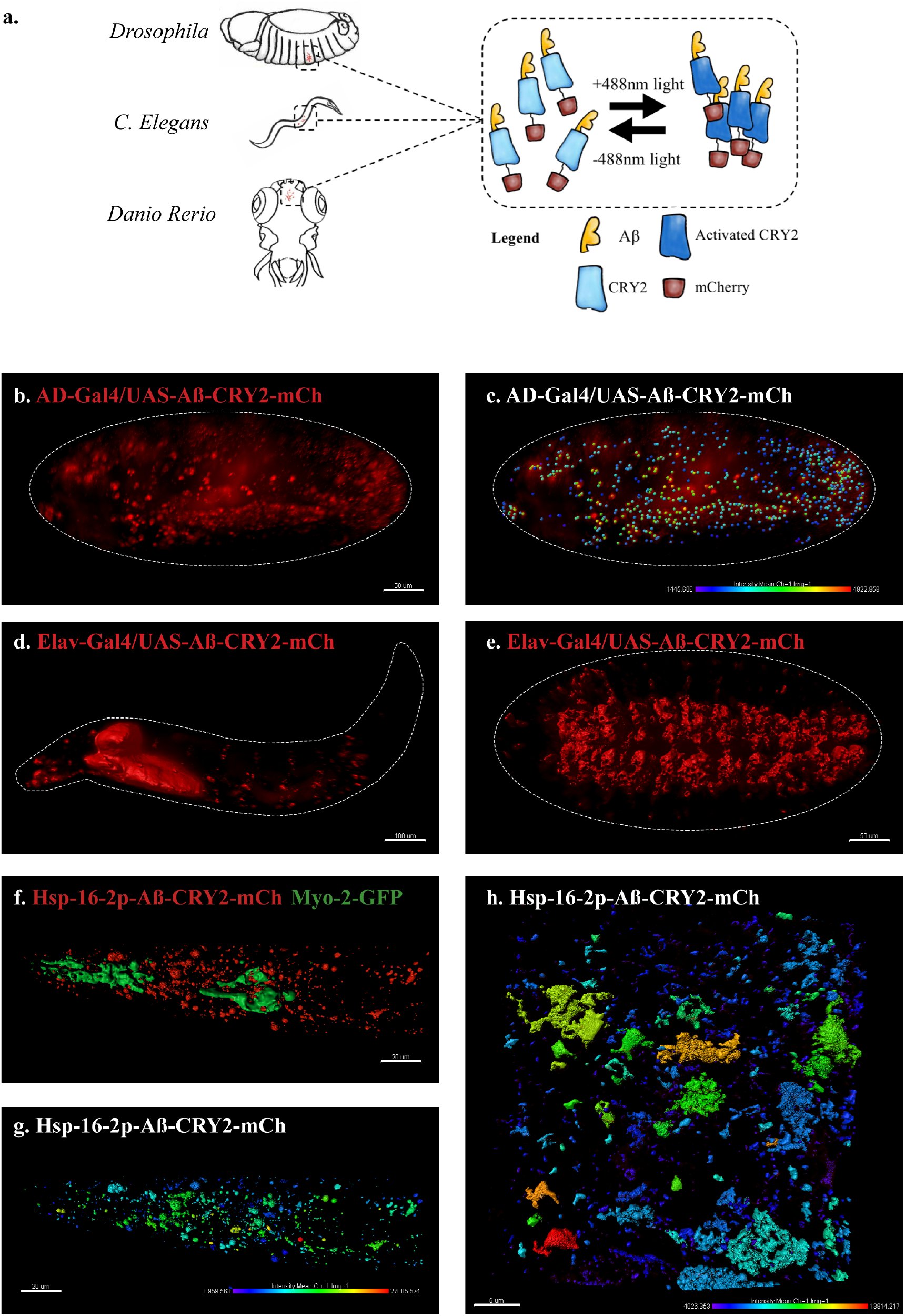
An *in vivo*, light-dependent, oligomerization switch for the formation or dissolution of Aβ aggregates in the fruit fly and nematode. **(a)** A schematic of the strategy. **(b)** Expression of AD-Gal4/UAS-Aβ-CRY2-mCh in a *Drosophila* embryo. **(c)** Mean intensity of aggregates in the same AD-Gal4/UAS-Aβ-CRY2-mCh *Drosophila* embryo. (**d & e**) Expression of Elav-Gal4/UAS-Aβ-CRY2-mCh in a *Drosophila* larva and embryo. **(f)** Expression of hsp-16-2p-Aβ-CRY2-mCh in heat-shocked *C. elegans* with *myo-2-gfp* marker, that marks its pharyngeal pump and serves as an indicator for positive microinjection. (**g & h**) Mean intensity of aggregates in the hsp-16-2p-Aβ-CRY2-mCh in *C.elegans* under 20× and 63× magnification of the confocal microscope; 63× image was processed using the Zeiss Airyscan.

### Generation of transgenic models with light-inducible intracellular Aβ oligomerization in model organisms

We generated transgenic animals expressing the 42-amino-acid human Aβ peptide (Aβ1-42) fused to Cryptochrome2 and the fluorescent protein mCherry (Aβ-CRY2-mCh). *Drosophila* lines were under the control of the GAL4/UAS system(Brand et al. 1993) and expressed either ubiquitously via AD-Gal4 driver (Figure 1b-c) or specifically in neurons using Elav-Gal4 driver (Figure 1d-e, S1). *C. elegans* lines were made with the heat shock promoter (Figure 1f-h, S2). In order to observe oligomerization and clustering of intracellular Aβ, we took a live imaging approach in *Drosophila* embryos using a lightsheet microscope with a 488nm laser activating CRY2 and the 561nm laser imaging mCh over time. We observed clusters of mCh fluorescence forming in embryos exposed to 488nm light, but fewer in embryos that were not exposed to blue light (Figure S3, Movies 1-2). The intensity of the clusters was quantified using Imaris quantification software to show the intensity of oligomer formation (Figure 1c).

We proceeded to test the effectiveness of light induced clustering of Aβ-CRY2-mCh in *C. elegans*. As this construct is driven by a heat shock promoter, worms were heat shocked at 35°C for 90 minutes followed by resting at 20°C before being immobilized and imaged using confocal microscopy (Figure 1f-h). We observed an increase in clusters in worms exposed to blue light over their siblings not exposed to blue light. The results were quantified as fluorescence intensity and represented in (Figure 1h) showing the formation of Aβ clusters. Even though the CRY2 system predominantly oligomerizes, we also observed the formation of intracellular Aβ aggregates, using a direct stain for Aβ aggregates (Figure S4). Importantly, we observed that Aβ aggregation was not reversible as had previously been observed for signaling molecules (Huang, Amourda et al. 2017, Johnson, Goyal et al. 2017, Kaur, Saunders et al. 2017), rather CRY2 appears to initiate clustering leading to Aβ bundles that do not come apart once blue light is turned off. We further confirmed that turning on blue light alone without the transgene expression did not lead to any Aβ expression or cluster formation (Figure S2b, Ab HS L vs Ab –HS L), suggesting that the oligomerization process is CRY2-specific and not an artifact of light. Together, our results showed that the Aβ-CRY2-mCh transgene is functional in both organisms, with a high specificity to blue light, which induces irreversible oligomerization of Aβ.

### Light-inducible Aβ oligomerization causes lifespan and behavioral deficits in transgenic models

To test the functionality of Aβ-CRY2-mCh, we next looked for phenotypes associated with light-induced intracellular Aβ oligomerization. We performed lifespan studies to assess the effects of Aβ-CRY2-mCh by expressing Aβ-CRY2-mCh either ubiquitously (AD-Gal4) or specifically in the nervous system (Elav-Gal4). Newly eclosed adult flies were separated into two groups in each case, one of which was exposed to ambient white light and the other kept in the dark. In both cases, flies exposed to light died much more quickly with half the mean and maximum lifespans (Figure 2a-b) compared to those reared in the dark. We extended this analysis to *C. elegans*, though starting at Day 6 due to the heat-shock manipulation step. Under light conditions, *C. elegans* showed markedly decreased lifespans as compared to the transgenic *C. elegans* kept in dark conditions (Figure 2c). Most significantly, we observed that the lifespan decrease was reversible, as taking animals exposed to light for 24 days and moving them into darkness showed a recovery of their lifespan, or a rescue (Figure 2a-b). To ensure that these reduced lifespans were physiological, we examined the fitness of the animals. Fitness was severely decreased in light-exposed flies and worms as quantified by fertility and locomotive assays (Figure S5). Interestingly, light-induced Aβ oligomerization caused further impairment in sensorimotor function of transgenic Aβ nematodes (Figure S5c). These results suggest that light-induced oligomerization of Aβ not only affected lifespan and fitness, but also impaired complex sensorimotor function related to neurobehavior.

**Figure 2.**
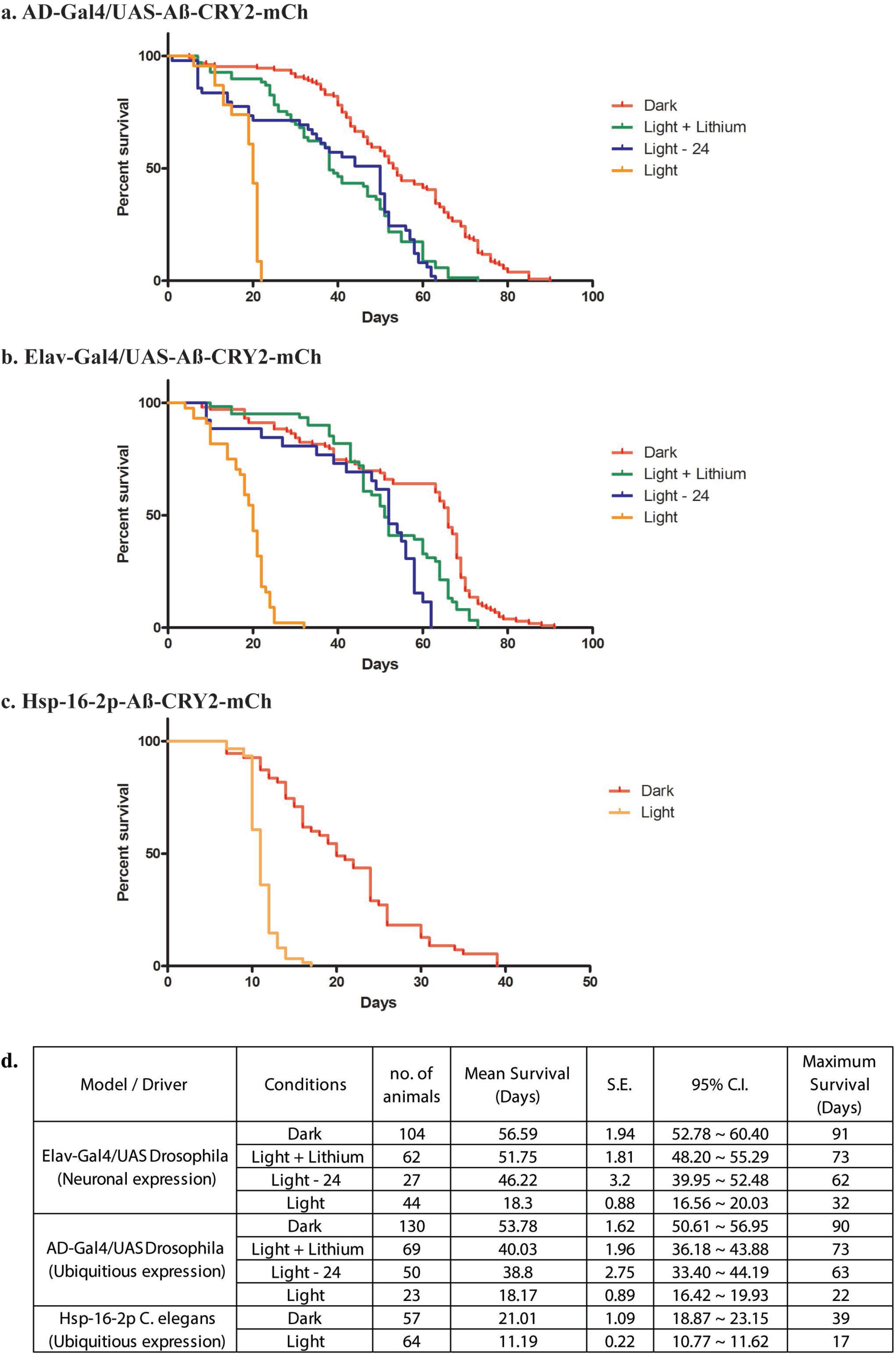
Analysis of lifespans in transgenic Aβ-aggregation *C. elegans* and *Drosophila* and adult models. **(a)** Kaplan-Meier survival curves of transgenic Aβ *Drosophila* adults driven by AD-Gal4 kept in the dark, exposed to light, exposed to light for 24 days then moved to the dark and exposed to light but fed with a lithium-supplemented diet. **(b)** Kaplan-Meier survival curves of transgenic Aβ *Drosophila* adults driven by Elav-Gal4 kept in the dark, exposed to light, exposed to light for 24 days then moved to the dark and exposed to light but fed with a lithium-supplemented diet. (c) Kaplan-Meier survival curves of control non-heat shocked nematodes, transgenic Aβ heat-shocked worms with the hsp-16-2p promoter kept in the dark and transgenic Aβ heat-shocked worms with the hsp-16-2p promoter kept in the light; those in light show a significantly shortened lifespan (overall significance by log-rank test: ****p < 0.0001) **(d)** Table showing number of samples used with the mean and maximum survival times for all conditions.

### Light-inducible Aβ oligomerization leads to physical and metabolic damage in transgenic models

The lethality mediated by light-induced Aβ oligomerization led us to examine the morphological effects of Aβ-CRY2-mCh clustering during embryonic stages in *Drosophila*. Embryos expressing Aβ in only the nervous system developed at a normal pace and did not show any morphogenetic defects and even hatched when in the absence of blue light (Movie 1). In contrast, sibling embryos exposed to blue laser light arrested in late neurogenesis stages where the central nervous system stopped developing and the embryos died (Movie 2 and Figure S3, S6). These embryos showed a variety of abnormalities in the positioning of neurons and glial cells (Compare to complete normal development, Movie 3) suggesting that light-induced Aβ oligomerization causes physical, structural damage that affects neuronal development.

The severity of the physical damage phenotype in embryos exposed to blue light (Figure 3a-d”) led us to investigate the differences in mechanism between the two conditions by exploring inflammatory response, mitochondrial health and oxidative damage. Expression of Aβ alone, without light-induced Aβ oligomerization, led to a small increase in inflammatory markers (Figure 3e)(Westfall et al. 2018). In embryos exposed to light, two of these markers increased to a much greater level (Figure 3e). We observed a significant reduction the number of mitochondria light-treated Aβ animals (Figure 3f). Markers of oxidative damage in light and dark exposed *C. elegans* showed differential expression of antioxidant defense genes, specifically Catalase, Trx-2 and Sod-3, suggestive of the occurrence of oxidative stress induced by Aβ oligomerization (Figure 4). These results suggested that light-induced Aβ oligomerization phenocopies many of the hallmarks of AD.

**Figure 3.**
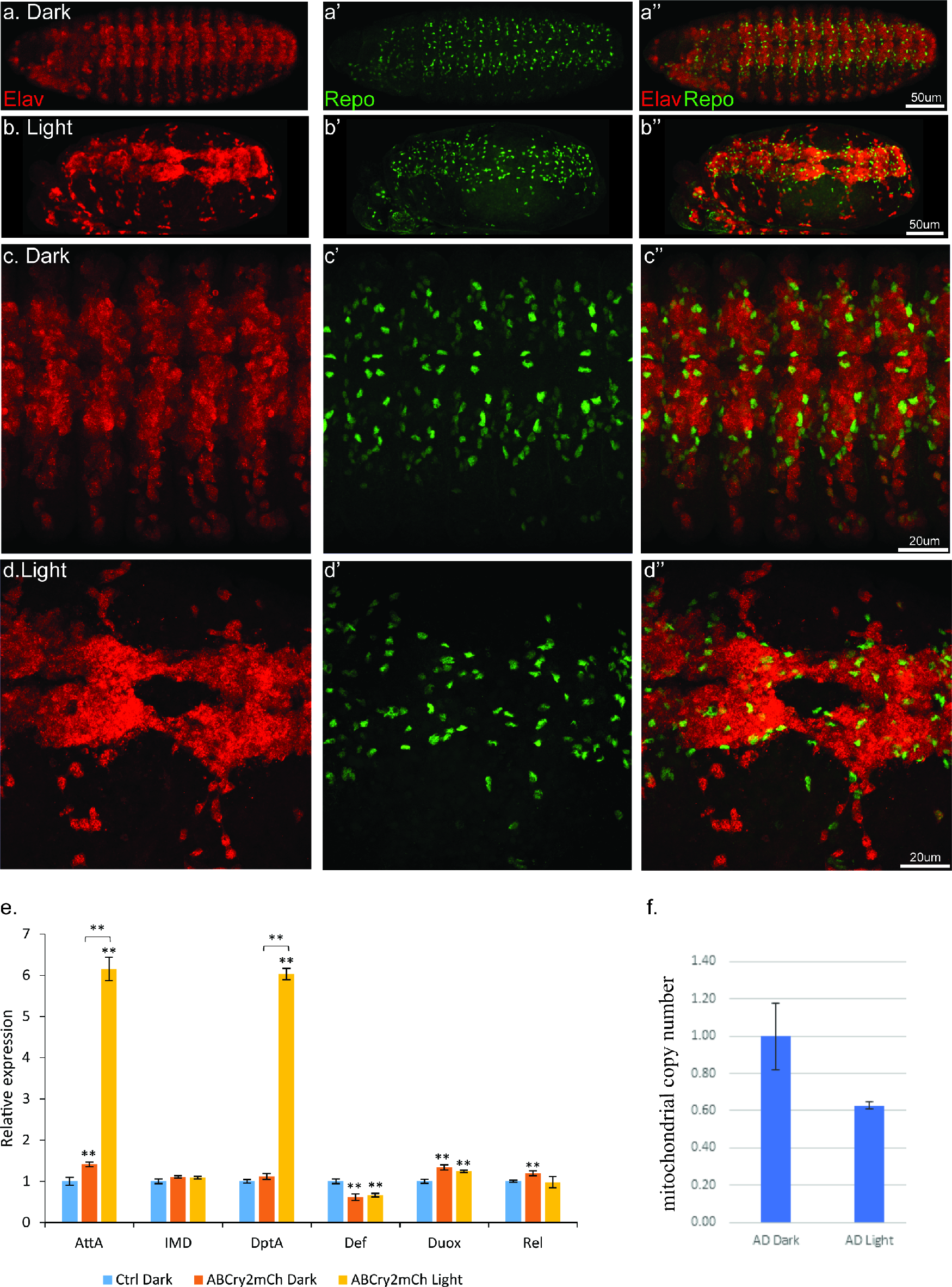
Light induced aggregation causes physical disruption of embryonic neural tissues. (**a-a”**) Elav-Gal4/UAS-Aβ-CRY2-mCh embryos kept in the dark and imaged for neurons (Elav) and glial cells (Repo). (**b-b”**) Elav-Gal4/UAS-Aβ-CRY2-mCh embryos exposed to light and imaged for neurons (Elav) and glial cells (Repo). (c-d”) Close ups. (E) Aggregation induced inflammatory response as measured by qPCR for inflammatory response genes. (f) The number of mitochondria in transgenic Aβ *Drosophila* embryo was reduced by 40% in light.

**Figure 4.**
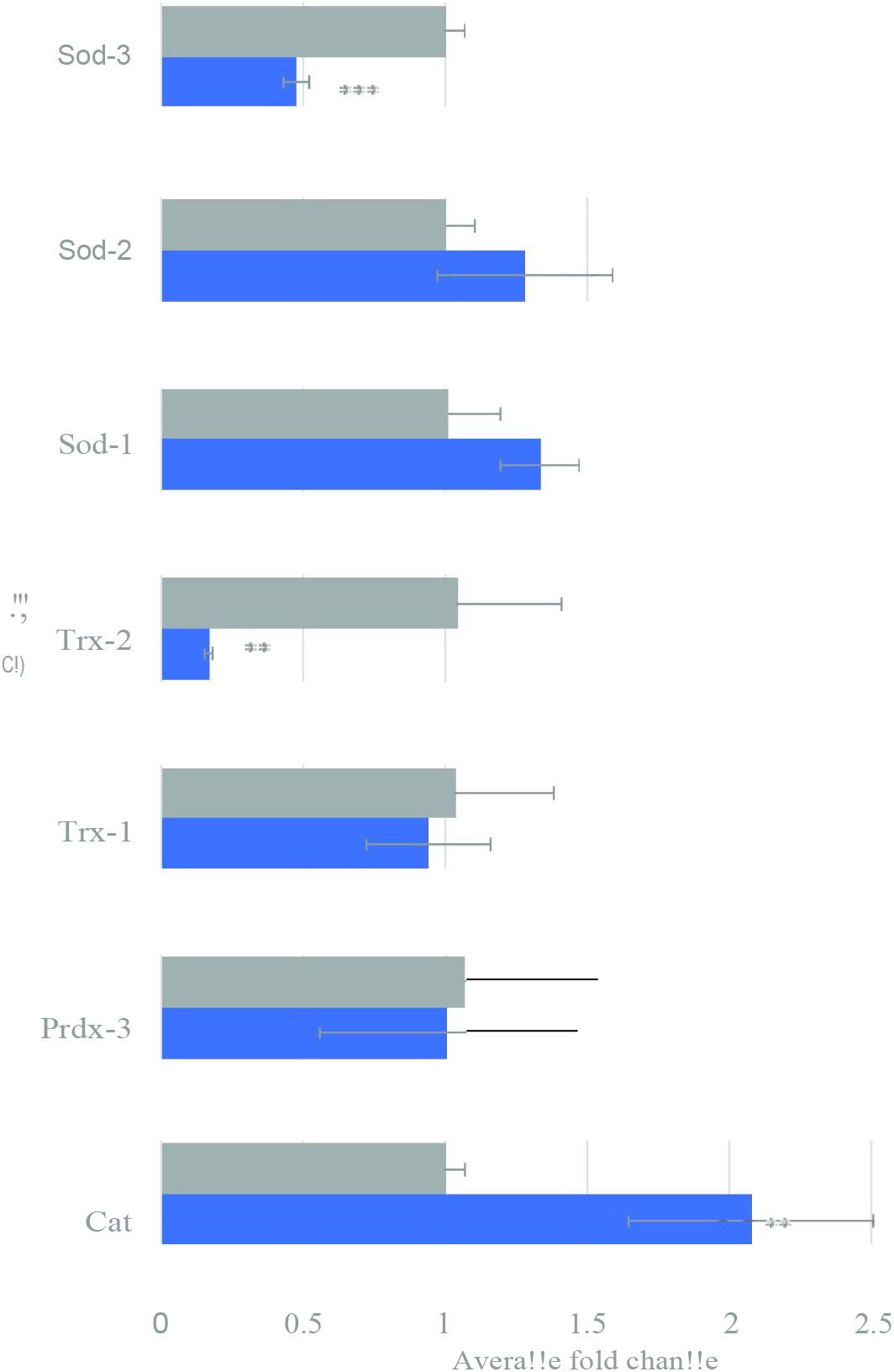
Gene expression of oxidative stress related genes in transgenic Aβ *C. elegans* with and without light-induction. Differential gene expression of catalase (Cat), peroxiredoxin-3 (Prd×3), thiroredoxin-1 (Trx-1), thioredoxin-2 (Trx-2), superoxide dismutase-1 (SOD-1), superoxide dismutase-2 (SOD-2) and superoxide dismutase-3 (SOD-3) in transgenic Aβ *C. elegan*s in light condition compared to control dark condition. Light-induced Aβ aggregation showed up-regulation in expression of Cat (**p <0.01) and down-regulation in expression of Trx-2 (*p <0.01) and Sod-3 (***p < 0.001).

We proceeded to look at light-induced Aβ oligomerization in metabolic impairments in transgenic *C. elegans*. Heat-shocked mutants exposed to light (Aβ HS L) had lower ATP levels compared to the non-transgenic control (Ctrl -HS D) (Figure 5). However, heat-shocked mutants in dark condition (Aβ HS D) did not show any significant differences in ATP level compared to the non-transgenic control. This result suggested that presence of Aβ alone is insufficient to induce ATP deficits, and that light-induced oligomerization of Aβ is required for the defect to manifest. Aβ expression alone was also insufficient to affect the nematodes’ maximum respiratory capacity, as heat-shocked mutants in dark condition (Aβ HS D) did not display differences in maximum and spare respiration capacity compared to control animals (Ctrl -HS D). However, light-induced Aβ oligomerization significantly reduced maximum respiration capacity in *C. elegans*. Heat-shocked mutants in the light condition (Aβ HS L) displayed significantly lower maximum and spare respiration capacity compared to controls (Ctrl -HS D) and to heat-shocked mutants in the dark condition (Aβ HS (Figure 5b-d). Together, these results highlight the metabolic differences between the two conditions and show that metabolic deficits are mediated by light-induced Aβ oligomerization.

**Figure 5.**
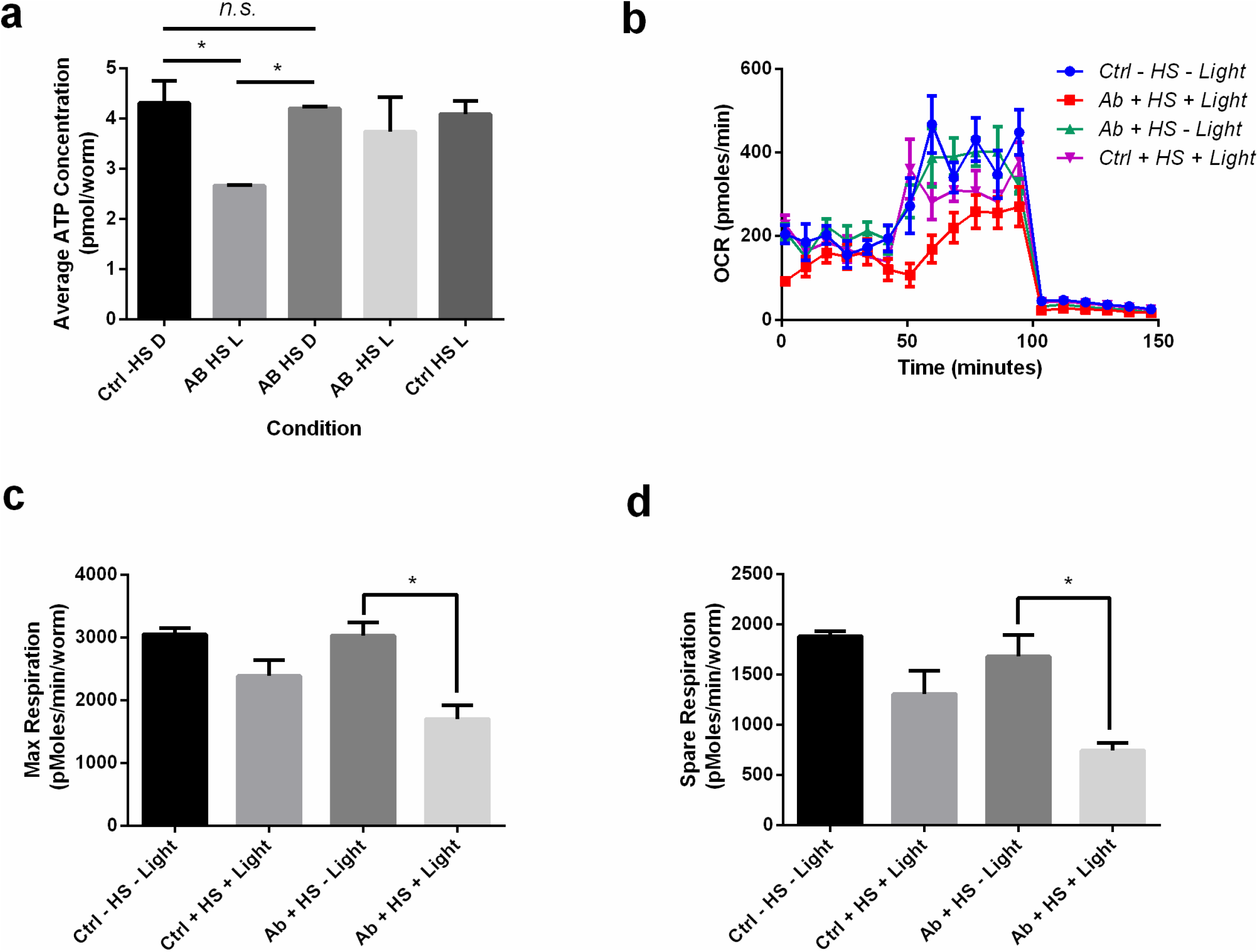
Light-induced Aβ oligomerization is required for the manifestation of metabolic defects in transgenic *C. elegans*. **(a)** ATP levels of control and transgenic Aβ nematodes (n=3 repeats for each condition, with 100 nematodes per repeat; ANOVA post-test p<0.05, *). **(b)** Oxygen consumption profile (OCR) of control and transgenic Aβ nematodes (n=6 repeats for each condition, with 10 nematodes per repeat). **(c)** Maximal respiration derived from OCR (n=6 repeats for each condition, with 10 nematodes per repeat; ANOVA post-test P<0.05, *). **(d)** Spare respiration derived from OCR (n=6 repeats for each condition, with 10 nematodes per repeat; ANOVA post-test p<0.05, *).

### Light-inducible Aβ lifespan decrease is rescued by Li^+^

Among the mechanisms for neuroprotection, the Wnt signaling pathway plays a role in modulating AD pathology and its progression (De Ferrari et al. 2003, Toledo et al. 2009, Parr et al. 2015, Jin et al. 2017). Activation of Wnt signaling through lithium (Li^+^) treatment attenuated Aβ aggregation and increased the lifespan of Aβ models (Toledo and Inestrosa 2009). By inhibiting the enzyme glycogen synthase kinase-3 (GSK-3), Li^+^ compounds drive the downstream production of intracellular β-catenin activating the Wnt–β-catenin signaling pathway(Klein et al. 1996). Li^+^ treatment through Wnt activation also rescued behavioral impairment and neurodegeneration induced by Aβ fibrils(De Ferrari, Chacón et al. 2003). Taken together, these studies point to a putative AD therapeutic intervention that is based on the activation of the Wnt signaling cascade through Li^+^. To test the usefulness of optogenetic Aβ for drug testing, we supplemented the food for *Drosophila* expressing Aβ-CRY2-mCh with Li^+^. *Drosophila* exposed to light, but consuming Li^+^ had significantly increased lifespans, suggesting a rescue effect induced by Li^+^ (Figure 2a-b), and that the optogenetic model can be used for inducible expression drug testing. Rapid drug testing can be also be achieved through cell culture. We extended this assay to HEK293 cells transfected with Aβ-CRY2-mCh. These showed Aβ clusters forming upon stimulation with blue light, but the clustering was ameliorated by the addition of Li^+^ or the specific GSK3 inhibitor CHIR99021 (Figure S7). These data demonstrate the potential use of our optogenetics system for drug testing.

Finally, we also extended these findings to a vertebrate system. We generated a zebrafish permanent line with UAS: Aβ-CRY2-mCh in the transgenic *TgBAC(gng8:GAL4) c416* background (Hong et al. 2013). We observed a diffuse mCh fluorescence in a small subset of neurons in the olfactory epithelium, the interpeduncular nucleus and in a few neurons sparsely distributed in the forebrain at 5 dpf. Upon blue light exposure (488 nM) the detectable intensity of the fluorescence increased rapidly (Figure S8). Several neurons among those expressing appeared to bleb and began to die within 40 to 50 minutes of initial exposure to the blue light (Movie 4).

## Discussion

To the best of our knowledge, this is the first study to demonstrate that induced oligomerization of intracellular Aβ mediates AD pathologies and that different pathologies (induced by Aβ oligomerization vs expression alone) are separable. Previous approaches lacked an inducible oligomerization tool and could only demonstrate Aβ oligomers’ toxicity through exogenous injection of Aβ oligomers (Mroczko, Groblewska et al. 2018). We use a tool to investigate the pathological effects of intracellular Aβ oligomerization in nematodes and flies, human kidney cells, and in the vertebrate model organism zebrafish (Figure S8, Movie 4). In all three systems, the CRY2 initiated clumping of Aβ that appeared to be irreversible likely due to the structure of the aggregates (Antzutkin et al. 2012). Consistent with existing literature showing that Aβ oligomers induce oxidative stress and inflammation(Butterfield et al. 2013, Forloni et al. 2018), we show that light-induced oligomerization of Aβ, but not Aβ expression alone, is necessary for the metabolic defects, loss of mitochondria and inflammation in *C. elegans* and *Drosophila*. Light-induced oligomerization also leads to physical damage of the nervous system, resembling the loss of brain tissue in AD (Figure S8). The embryonic assay for physical damage phenocopies brain lesions only to some extent as the extent of the damage is enhanced by the forces applied to the developing neural tissue. The irreversibility of Aβ aggregates once initiated as seen here, also hints at intrinsic properties that may be unique or unusual to this peptide and warrants further investigation. It is also likely to prove a good model for testing anti-aggregation compounds.

We anticipate that our approach will serve as an attractive tool for carrying out drug screens and mechanistic studies for the treatment of AD. This optogenetic strategy will also undoubtedly complement other techniques due to the high level of spatiotemporal specificity. For example, it could be optimally positioned to gain insights into the unresolved mechanism of Aβ – whether oligomerization and subsequent accumulation or lack of effective clearance underpins AD pathologies. The separation of two phases of AD progression also suggests that single drug treatments will not suffice, and perhaps combinatorial approaches should be tried.

## Materials and Methods

### Molecular Cloning of Optogenetic Transgenes

DNA sequences corresponding to the human Amyloidβ1-42 amino acid sequence were synthesized (IDT) into a Gateway technology (Invitrogen) entry vector. The human Aβ1-42 sequence was used for *Drosophila* and zebrafish due to close evolutionary conservation and a nematode-codon-optimized version of Aβ1-42 was used for *C. elegans* (Fong et al. 2016). The CRY2-mCh fragment was cloned into pDONR vector with attB5 and attB2 sites, and MultiSite Gateway® Pro 2.0 recombination (Invitrogen) was used to recombine donor plasmids and pDONR-CRY2-mCh into the respective destination vectors for three model organisms:

a. pUASg.attB.3XHA for *Drosophila* to synthesize pUAS-Aβ-CRY2-mCh (Bischof et al. 2007)
b. pDEST-hsp-16-2p, a kind gift from Hidehito Kuroyanagi (Medical Research Institute, Tokyo Medical and Dental University) to synthesize hsp-16-2p-Aβ-CRY2-mCh for *C. elegans* (Okazaki et al. 2012).
c. pDEST40 (Invitrogen) was used to make CMV driven pDEST40-Aβ-CRY2-mCh for expression in *D. Rerio* and HEK293 tissue culture cells.

### Crosses and expression of UAS construct

For *Drosophila*, the transgenes were injected into attP2 (Strain#8622) P[CaryP]attP268A4 by BestGene Inc. (California)(Groth et al. 2004, Markstein et al. 2008). Expression was driven by Elav-GAL4 the neuronal driver and two ubiquitous drivers armadillo-GAL4 and daughterless-GAL4 (Brand and Perrimon 1993). All additional stocks were obtained from the Bloomington Drosophila Stock Center (NIH P40OD018537) were used in this study.

Fly crosses performed were:

1. Elav-Gal4 × w; UAS-Aβ-CRY2-mCh
2. Arm-Gal4; Da-Gal4 × w; UAS-Aβ-CRY2-mCh
3. Elav-Gal4; Repo-QF2, QUAS-GFP × UAS-Aβ-CRY2-mCh
4. Elav-Gal4; Repo-QF2, QUAS-GFP × UAS-tdTomato

For *C. elegans*, 50ng/μl the construct hsp-16-2p-Aβ-CRY2-mCh was co-injected with 25ng/μl of pharyngeal-specific fluorescence marker *myo-2-gfp* into the distal gonads of wild type young adults. A *myo-2-gfp* strain *C. elegans* was also generated and used alongside as controls.

For *D. rerio*, 70ng/μl of the construct pDEST40-Aβ-CRY2-mCh was injected to zebrafish embryos during the one-cell stage. Wild type and nacre strains were used. Potassium chloride was used as the control for microinjection.

### Animal husbandry

*Drosophila* were maintained at standard humidity and temperature (25°C) with food containing 6g Bacto agar, 114g glucose, 56g cornmeal, 25g Brewer’s yeast and 20ml of 10% Nipagin in 1L final volume. Transgenic *Drosophila* and controls were distributed into either dark or light condition on Day 1. *C. elegans* were maintained as previously described at 20°C(Stiernagle). Age-synchronized nematodes were generated by hypochlorite bleaching. 250μM of 5-fluoro-2’-deoxyuridine (FUDR) was added to prevent progeny production in all experiments except for fertility assays. For Fertility assay, normal Nematode Growth Media (NGM) agar plates were used. To induce expression of Aβ-CRY2-mCh, Day 4 young adult worms were heat shocked at 35°C for 90 minutes, and subsequently incubated at 20°C for a day (Day 5) for recovery. The heat-shocked transgenic Day 6 *C. elegans* were then separated into dark or light condition and was maintained at 20°C throughout the experimental period. Non-heat shock controls were used for lifespan, fertility and locomotion studies.

### Lifespan Assays

*Drosophila* and *C. elegans* were counted daily for the number of dead subjects and the number of censored subjects (excluded from the study). *Drosophila* that failed to respond to taps were scored as dead and those stuck to the food were censored. *C. elegans* that failed to respond to plate-tapping were scored as dead, those that burrowed to the side of the plate were censored.

### Locomotion assay

*Drosophila* locomotion was assessed using an established geotaxis assay previous described in(Rival et al. 2009). In brief, 25 *Drosophila* were enclosed in a plastic column (25cm tall, with a 1.5cm internal diameter) and were tapped to the bottom. The number of *Drosophila* at the top (Ntop) of the column, and that at the bottom (Nbot) were counted after 20 seconds. Three trials with the same sample were performed within 30 seconds interval. Performance index was defined as (15+Ntop-Nbot)/30.

Heat-shocked *C. elegans* were assessed at Day 9, by placing worms onto individual NGM plates, pre-spotted with *Escherichia coli* OP50. *C. elegans* were placed on the side of the bacteria spot on the NGM plate using a platinum worm picker and were left undisturbed to move freely for 15 minutes. Distance travelled was determined by imaging and measuring of worm tracks on bacteria using a stereomicroscope.

### Fertility assay

25 pairs of male & female *Drosophila* expressing MD-Aβ-Cry2-mCh were kept in a single tube under dark or light conditions. The number of eggs laid were scored after 5 hours 30 minutes for Days 5, 6, 7, 13, 14 and 20. Day 5 heat-shocked *C. elegans* were transferred onto individual NGM plates spotted with *E. Coli*. Each worm was transferred to a fresh plate daily from Day 6 to Day 8, allowing 24 hours’ egg laying period. The number of new young worms were counted on each plate as the progeny of a single individual.

### Food race assay

Food race assay adapted from Nyamsuren *et al.(Nyamsuren et al. 2007)* was conducted using 94mm NGM plates seeded with 0.2mL of OP50-1 at a designated food spot. A starting spot 60mm away from the food spot was marked on the plate. 30 D2 adult worms from each condition 24hrs post heat-shock were placed on the starting spot and left to roam at room temperature. After 15min, the number of worms that reached the food spot and the number of worms that left the starting spot were tabulated. Mobility index was calculated by dividing the number of worms that reached the food spot by the number of worms that left the starting spot. 3 technical repeats were conducted for each condition.

### Adenosine triphosphate (ATP) assay

ATP assay using firefly lantern extract (Sigma-Aldrich) was performed as described by Tsujimoto(Tsujimoto et al. 1970). 100 D2 adult worms from each condition 24hr post heat-shock were freeze-thawed in liquid nitrogen and sonicated in 10% Trichloroacetic acid (TCA) buffer. The extracted ATP from the different conditions and ATP standards were loaded onto a 96-well plate and injected with firefly lantern extract. Luminescence was measured using a Cytation 3 Imaging Reader (Biotek).

### Mitochondrial metabolic flux assay

Mitochondrial metabolic flux assay was performed as described by Fong et al.(Fong et al. 2017). 60 D2 adult worms from each condition 24hr post heat-shock were transferred to a 96-well Seahorse plate containing M9 buffer. The plate was loaded into the XF96 Extracellular Flux Analyzer (Seahorse) which measures the animal’s basal and maximum respiratory capacity.

### Light-sheet microscopy

*Drosophila* embryos were dechorionated using bleach, rinsed twice with water and dried, and loaded into a capillary filled with 1% low-melting agarose Type VII-A in water (Sigma) (Colosimo et al. 2006). *C.elegans* and *D. rerio* were embedded directly in the agarose. Samples were imaged with a Lightsheet Z.1 microscope. Cry2 was activated by a dual-side illumination with 10% power 488nm laser for 29.95ms for every 2.5minutes for 500 cycles. Controls were only exposed to 561nm. Images were acquired with a water immersion objective at 10×/0.2 Illumination Optics and W Plan-Apochromat 20×.1.0 UV-VIS detection objective (Carl Zeiss, Germany). Image data were processed using the maximum intensity projection function of ZEN 2014 SP software (Carl Zeiss, Germany), and were analyzed with ImageJ (NIH) and IMARIS 9.0 (Bitplane AG, UK).

### Transfection and Live cell confocal microscopy

Human Embryonic Kidney (HEK) cells were obtained from ATCC and maintained in Dulbecco’s Modified Eagle’s Medium with 10% Fetal Bovine Serum and 1% Penicillin and Streptomycin (Invitrogen). Cells were plated at 80% confluence on 35mm TC treated glass bottom dish 24 hours prior to transfection with 2.5mg of pDEST40-Aβ-CRY2-mCh using the Effectene Transfection Reagent (Qiagen). Untransfected cells were used as negative control. Cells were imaged 24 hours after transfection using the Zeiss LSM800 confocal microscope with 63x oil immersion lens.

### Immunofluorescence & Confocal microscopy

*Drosophila* embryos 9 hours after deposition were incubated at 25°C in either light or dark conditions for 13 hours before being dechorionated using bleach. Embryos were then fixed with Heat-Methanol treatment (Müller et al. 1996) or with heptane/4% formaldehyde in phosphate buffer (0.1M NaPO4 pH 7.4) (Tolwinski et al. 2001). Staining, detection and image processing as described in(Colosimo and Tolwinski 2006).

Primary antibodies used were the glial cell marker anti-Repo (mouse mAb, 8D12) and the neuronal cell marker anti-Elav (rat mAb, 7EA810) from Developmental Studies Hybridoma Bank (DSHB developed under the auspices of the NICHD and maintained by The University of Iowa, Department of Biological Sciences, Iowa City, IA 52242). Secondary antibodies used were Alexa Flour 488 anti-rat and Alexa Flour 647 anti-mouse (Invitrogen).

For detection of Aβ aggregates, thioflavin T (ThT) staining were performed as previously described (Iijima et al. 2008). Formaldehyde-fixed embryos were incubated in 50% EtOH containing 0.1% ThT (Sigma) overnight. Embryos were destained in 50% EtOH for 10 minutes, followed by three washes in PBS. Embryos were then mounted on microscope slides using Aquapolymount (Polysciences, Inc.).

Images were acquired on the Zeiss LSM 800 (Carl Zeiss, Germany) using the following settings: 1% laser power for 488nm; 5% laser power for 561nm; 2% laser power for 647nm. Images were processed using the ZEN 2014 SP1 software (Carl Zeiss, Germany) and Imaris (Bitplane AG).

### RNA Extraction, cDNA Synthesis and qPCR

*Drosophila* embryos 9 hours after deposition were incubated at 25°C in either light or dark conditions for 13 hours were used. Embryos were dechorionated and washed with 100% ethanol prior to RNA extraction using the ISOLATE II RNA Mini Kit’s protocol (Bioline, UK). The extracted RNA was quantified using Nanodrop (Thermo Fisher Scientific). cDNA synthesis was done according to the SensiFAST™ cDNA Synthesis Kit‘s protocol (Bioline). Primers pairs used: AttA, IMD, DptA, Def, Duox, Rel, catalase, Prdx3, Trx1, Trx2, SOD-1, SOD-2, SOD-3 and reference gene Rpl32. Quantitative PCR was performed using SYBR® Green. Expression data were normalized to the dark controls. For mitochondrial copy number, the above collection method was used except the following: the embryos were stored in tritonX-100 and the primers used were Mitochondria Cytochrome b (MtCyb) and reference gene RNAse P.

### Image Data & Statistical Analysis

Aβ aggregates were quantified in *Drosophila* using ImageJ (NIH) and in *C. elegans* with IMARIS. For statistical analysis of the expression of genes, student’s t-test was used for except for cases where the data showed unequal standard deviation (F-test, p<0.05), in which the Mann-Whitney nonparametric test was performed. Statistical analysis of lifespan studies and behavioral assays were performed using OASIS2 6. The number of samples was determined empirically. All graphs were plotted using Graphpad PRISM 6 (Graphpad Software).

## Movies

**Movie 1:** Elav-Gal4/UAS-Aβ-CRY2-mCh embryos kept in the dark and imaged for neurons in red mCherry and glial cells Repo-QF2>QUAS-GFP. Blue laser power kept low to allow imaging of glial cells, but not high enough to activate aggregation.

**Movie 2:** Elav-Gal4/UAS-Aβ-CRY2-mCh embryos exposed to light and imaged for neurons in red mCherry and glial cells Repo-QF2>QUAS-GFP. Blue laser power at higher setting to allow imaging of glial cells and activate aggregation.

**Movie 3:** Elav-Gal4/UAS-MyrTomato embryos exposed to light and imaged for neurons in red (tdTom) and glial cells Repo-QF2>QUAS-GFP. Blue laser power at higher setting to allow imaging of glial cells and control for laser damage.

**Movie 4:** Expression of Aβ-CRY2-mCh in zebrafish developing brains. Schematic of imaging set up and area of interest. Imaging of cell movement, Aβ-CRY2-mCh aggregation and cell death.

---

**Figure S1.**
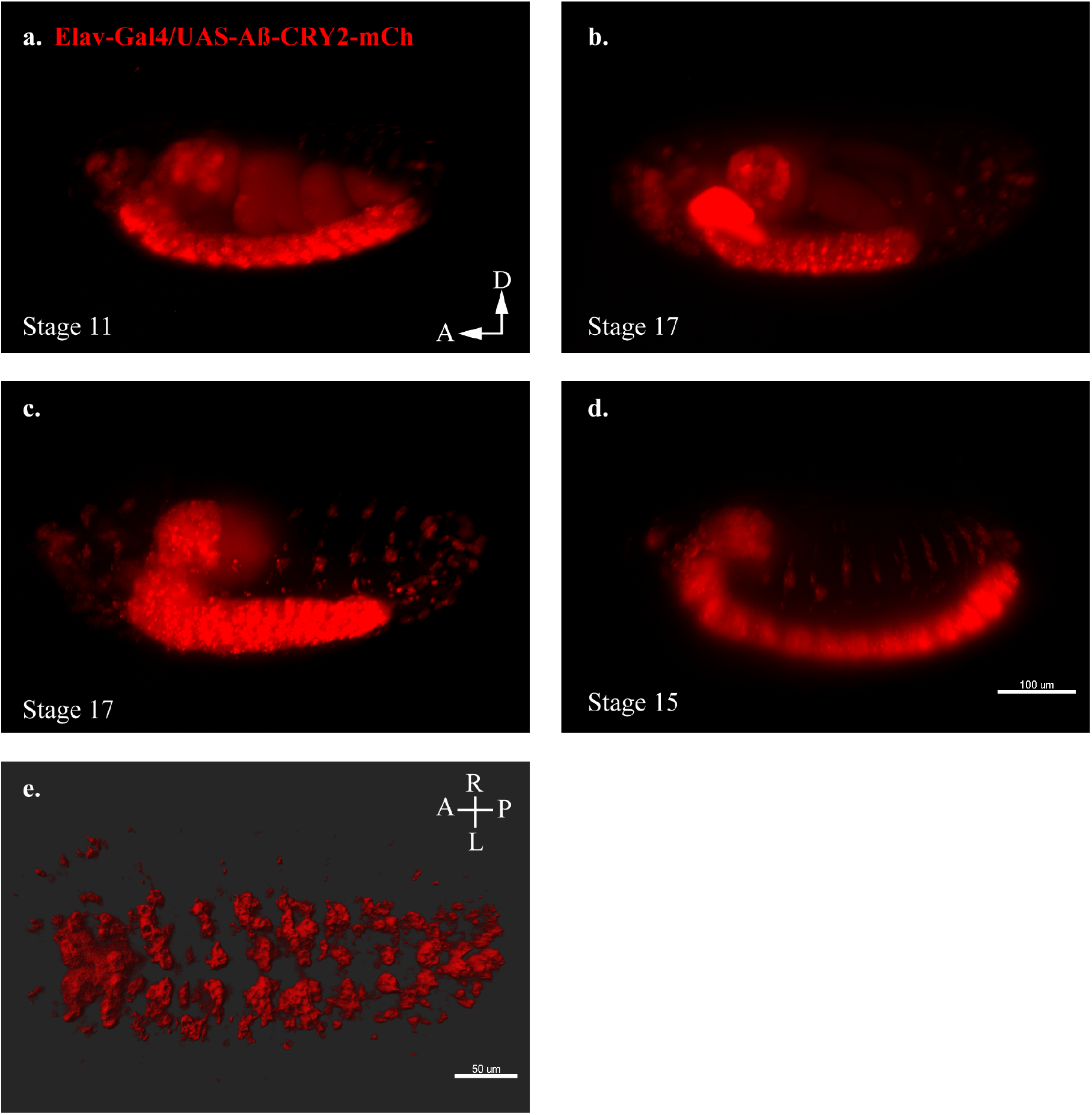
Transgenic *Drosophila* model expressing Aβ-CRY2-mCh. **(a-d)** Still images from lightsheet timelapse microscopy of Elav-Gal4/UAS-Aβ-CRY2-mCh in neurons detected by mCh fluorescence (red) in transgenic *Drosophila* embryos at different developmental stages. **(e)** Aβ localization to neuronal cell body volume in a *Drosophila* embryo expressing the mentioned construct.

**Figure S2.**
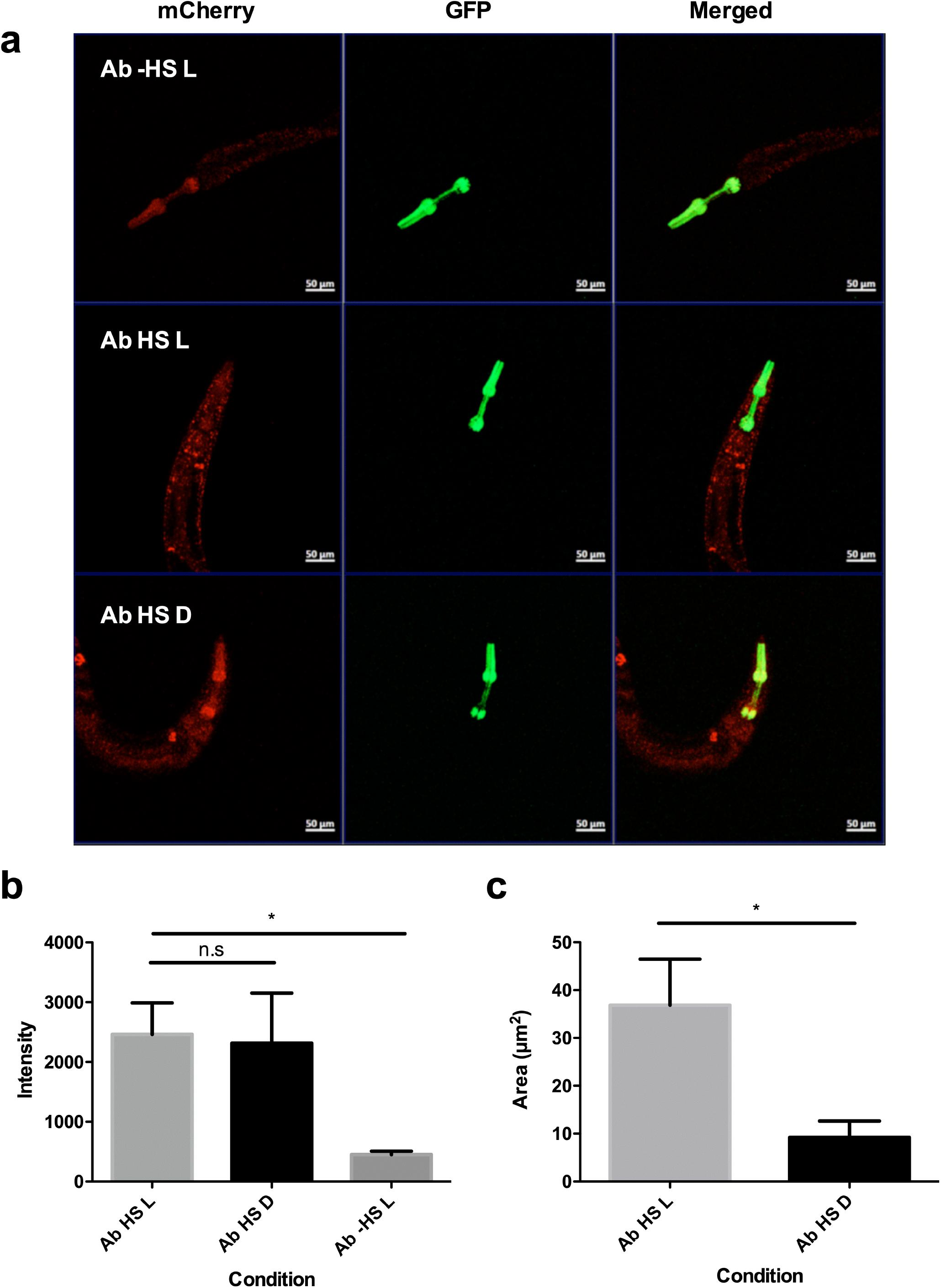
Exposure to light drives Aβ aggregation in *C. elegans* strains 24h post heatshock. **(a)** Confocal microscopy images of mutant *C. elegans* 24 hours post heat-shock at 20× magnification in Aβ -HS L (no heat shock, exposed to light), Aβ HS L (with heat shock, exposed to light) and Aβ HS D (with heat shock, kept in the dark) conditions. Aβ -HS L worms are used as a control for quantifying baseline Aβ expression. Distinct puncta (yellow) can be seen in animals in Aβ HS L condition. No red fluorescence was observed in control animals without the transgene (not shown). **(b)** Quantification of intensity of red fluorescence measured in the animals in Aβ HS L (n = 13), Aβ HS D (n = 9) and Aβ -HS L (n = 5) conditions 24 hours post heat-shock. Difference in expression between Aβ HS L conditions and Aβ -HS L is statistically significant (p = 0.034). (C) Quantification of total area of bright puncta in the animals in Aβ HS L (n = 13), and Aβ HS D (n = 9) conditions 24 hours post heat-shock. Difference in area of fluorescence between Aβ HS L conditions and Aβ HS D is statistically significant (p = 0.031).

**Figure S3.**
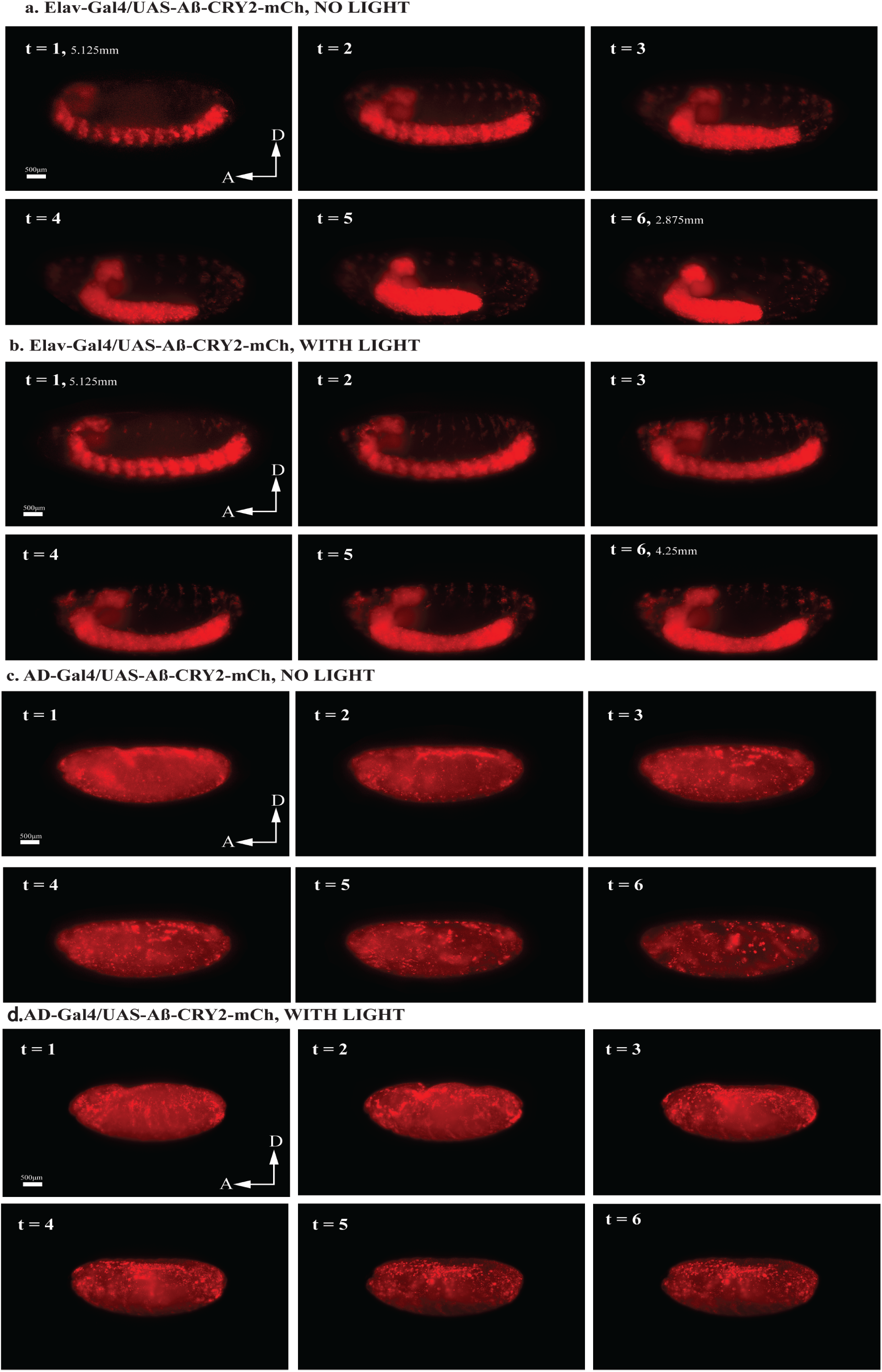
Light-induced Aβ aggregation in *Drosophila* embryos resulted in distinct developmental defects. **(a)** Still images from a 24-hours recording of Elav-Gal4/UAS-Aβ-CRY2-mCh aggregates in a *Drosophila* embryo without blue light illumination resulted in normal development. Time stages t=1-6 showed embryonic development stages 12-13 (~440-620 minutes after fertilization), where germ band retraction starts and ends respectively. Germ band was 5.125mm at t=1, and retracted to 2.875mm by t=6. **(b)** Still images from a 24-hours recording of Elav-Gal4/UAS-Aβ-CRY2-mCh aggregates in a *Drosophila* embryo with blue light illumination every 2.5 minutes resulted in the arrest of embryogenesis during late germ band retraction stages. Time stages t=1-6 showed embryonic development from late stage 10 to early stage 12 (~300-460 minutes after fertilization), where germ band retraction begins (stage 12). Germ band was 5.125mm at t=1, and retracted only to 4.25mm by t=6. **(c)** Still images from a 24-hours recording of uniform expression AD-Gal4/UAS-Aβ-CRY2-mCh aggregates in a *Drosophila* embryo without blue light illumination resulted in normal development. Time stages t=1-6 showed embryonic development stages 14-16 (~620-900 minutes after fertilization), where dorsal closure of midgut and epidermis and shortening of ventral nerve cord occurs. In this case, the ventral nerve cord shortening was not clearly visible. **(d)** Still images from a 24-hours recording of AD-Gal4/UAS-Aβ-CRY2-mCh aggregates in a *Drosophila* embryo with blue light illumination at every 2.5 minutes resulted in the arrest of embryogenesis at dorsal closure stages. Time stages t=1-6 corresponded to embryonic development stage 14 (approximately 620 – 680 minutes after fertilization), where dorsal closure begins.

**Figure S4.**
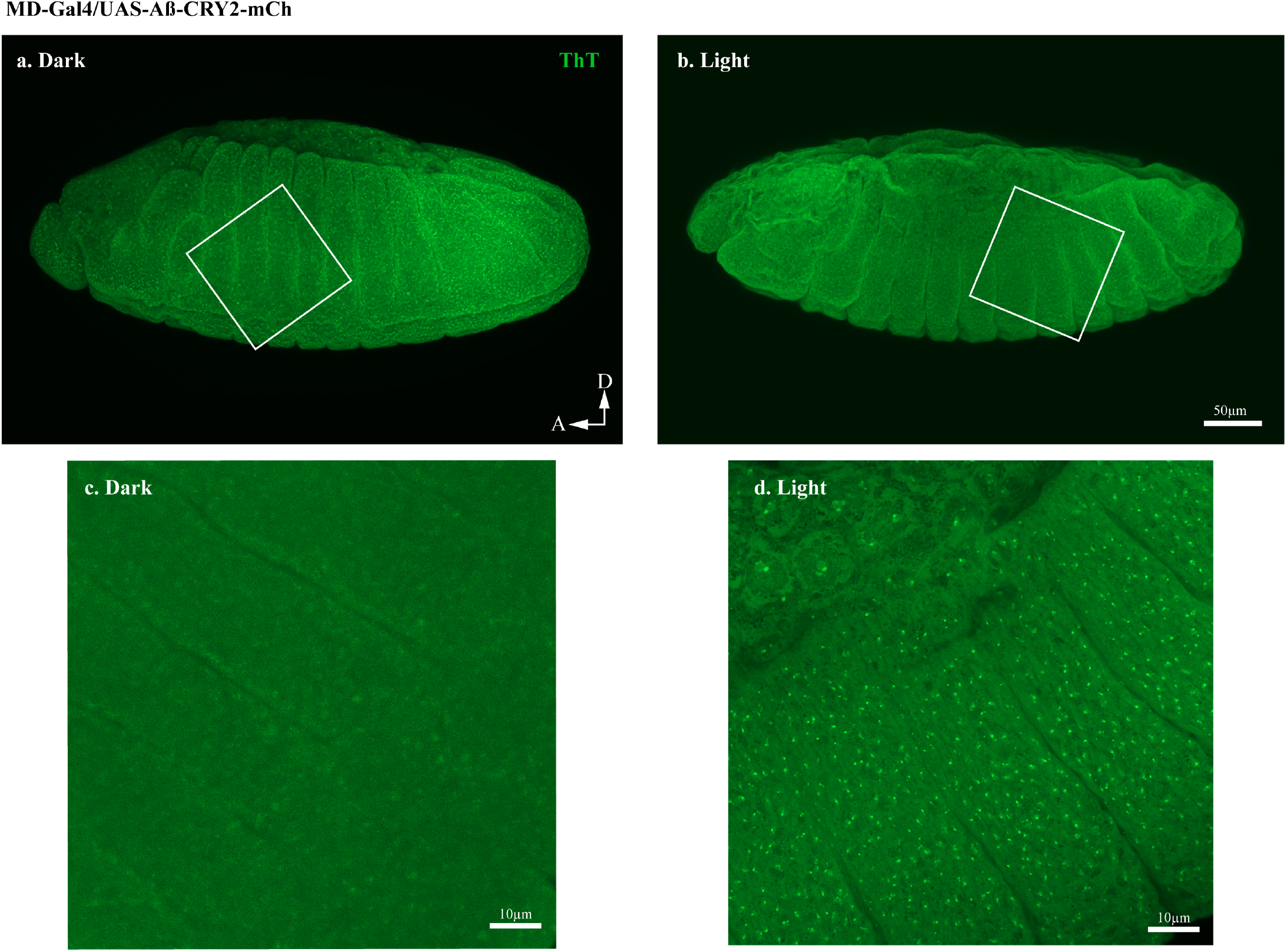
Thioflavin T (ThT) staining showing the Aβ aggregation transgenic *Drosophila* embryos ubiquitously expressing Aα-CRY2-mCh. Embryos expressing A²-CRY2-mCh ubiquitously showed significantly lesser Aâ aggregates in the (a) dark as compared to (b) embryo exposed to light. (a, b) were imaged at 20×. (c, d) were magnified images of the box in (a, b) imaged at 63×.

**Figure S5.**
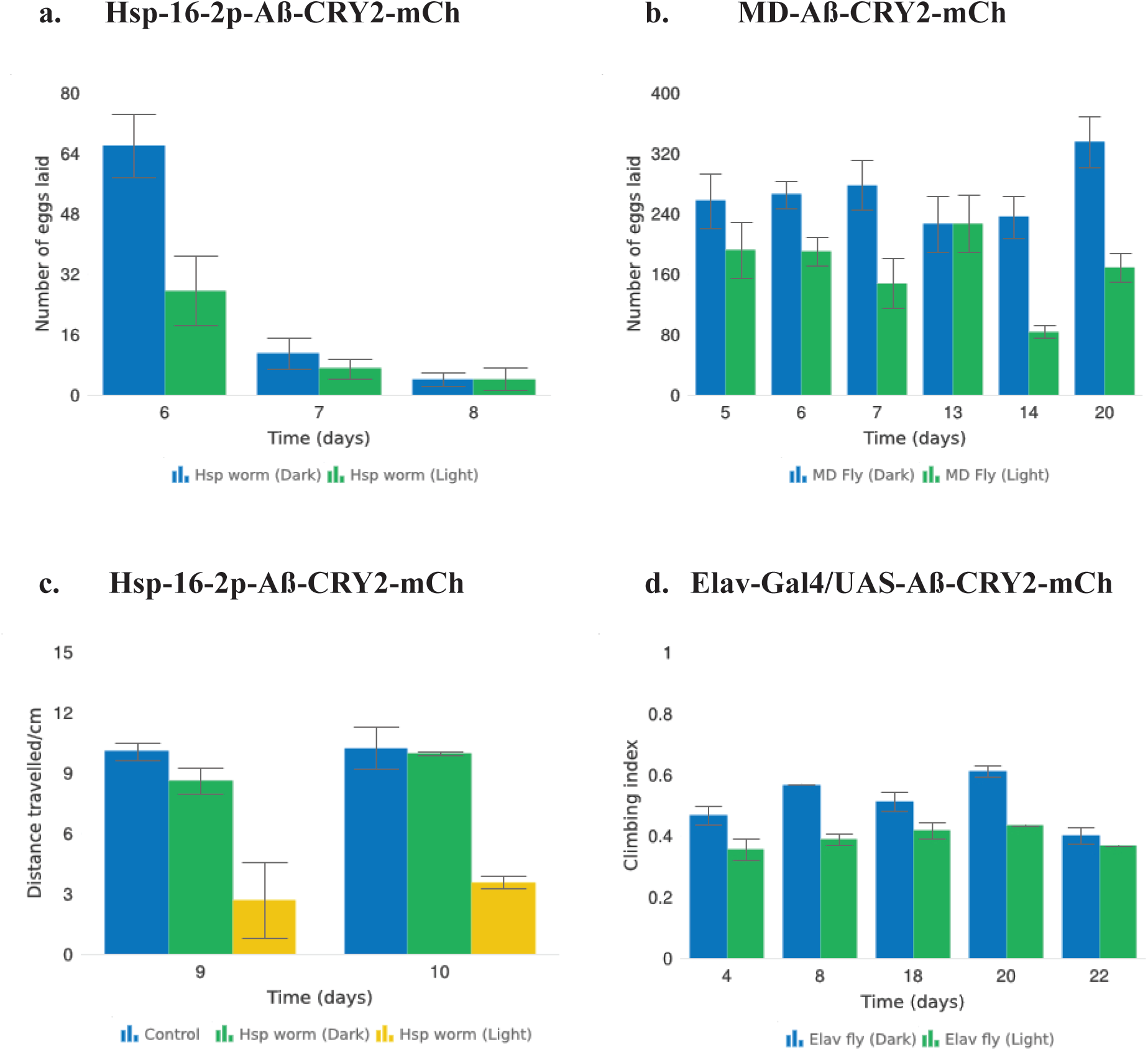
Aβ-CRY2-mCh expressing transgenic *C. elegans* and *Drosophila* reared in light showed signs of reduced fitness. **(a)** Exposing Aβ-CRY2-mCh *C. elegans* to blue light significantly reduced progeny numbers. Two-way ANOVA, p = 0.1. Data were sampled over three days (Days 6-8), n=8 light and dark worms. **(b)** MD-Gal4 driven Aβ-CRY2-mCh transgenic *Drosophila* produced significantly lower number of progeny when exposed to blue light. Two-way ANOVA ****p < 0.0001. Data were sampled on Days 5, 6, 7, 13, 14, and 20, n= 25 light and dark pairs of *Drosophila*. **(c)** Aβ-CRY2-mCh *C. elegans* have significantly reduced locomotive ability when reared in light. Distance travelled was reduced by about three-folds compared to the dark control and non-heat shocked control. Data were sampled on Days 9 and 10, two-way ANOVA **p < 0.01 n= 6 light, dark and non-heat shocked controls. **(d)** Climbing assay showed that transgenic Aβ *Drosophila* displayed sensorimotor impairments when exposed to light. Two-way ANOVA ****p < 0.0001. n= 18 light and dark conditions.

**Figure S6:**
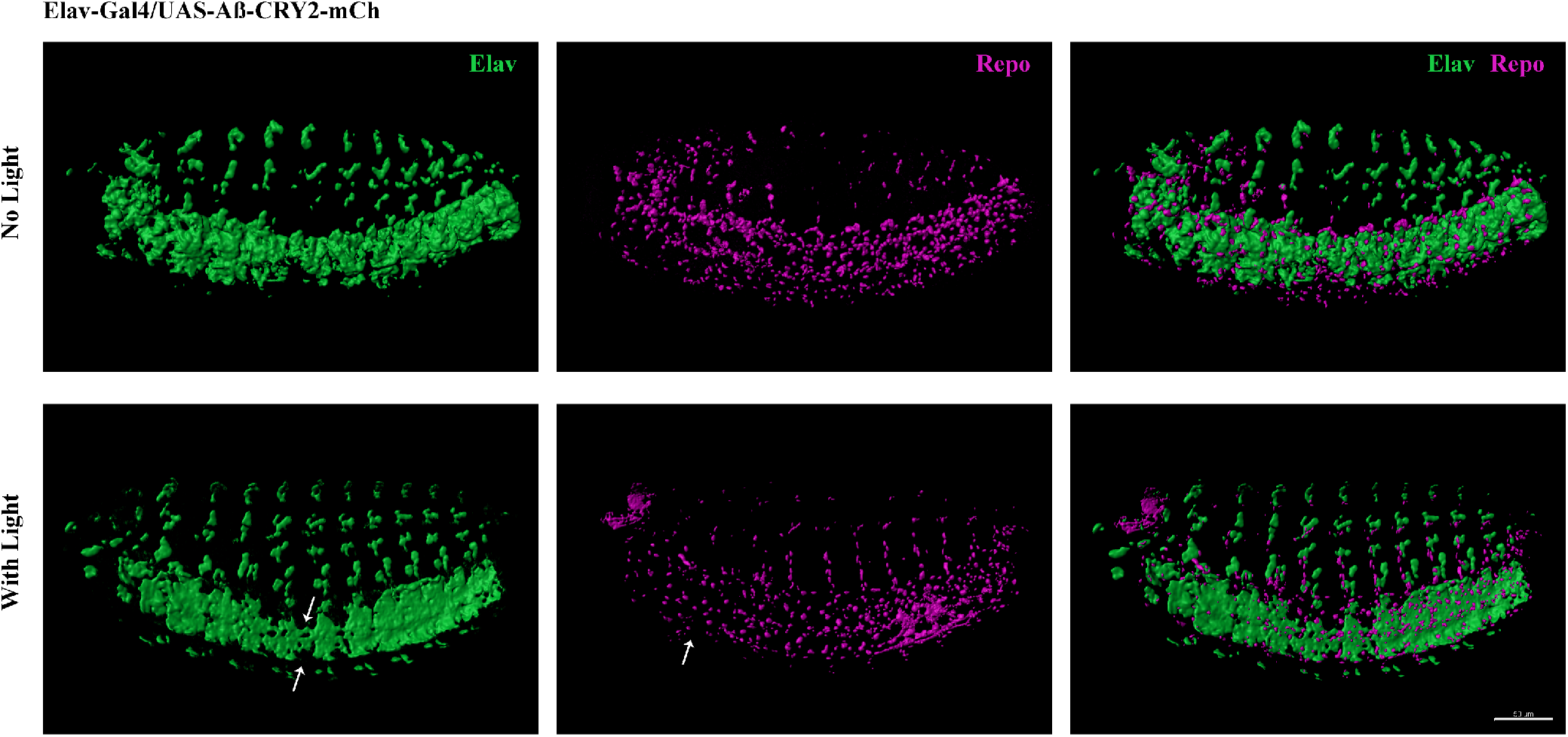
Light-induced Aβ aggregation leads to developmental deficits in transgenic *Drosophila* embryo. Reduction of neurons and glia cells (arrows) detected by immunostaining of Elav and Repo in transgenic Aβ *Drosophila* embryos imaged at 40×

**Figure S7.**
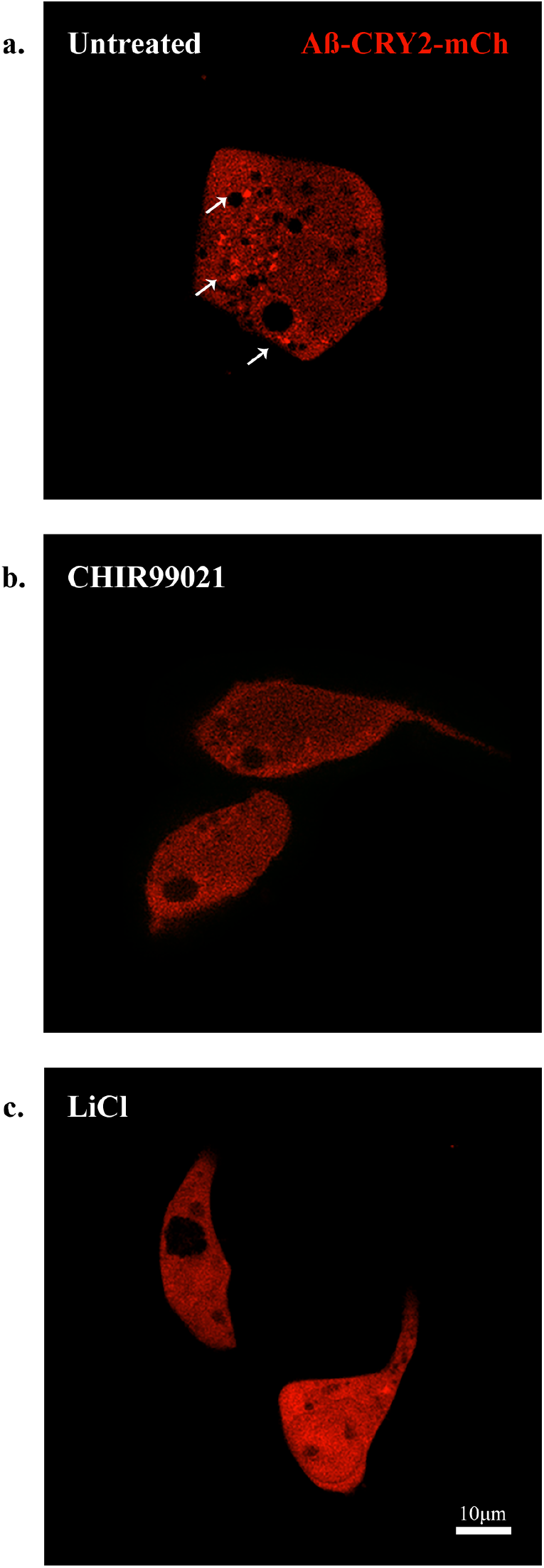
HEK cells expressing Aβ-CRY2-mCh in light condition. **(a)** Untreated, negative control HEK cells show light-induced clustering of Aβ-CRY2-mCh (white arrows) and increased number of vacuoles under blue light illumination. **(b)** HEK cells treated with GSK3 inhibitor CHIR99021 (positive control) show little clustering of the light-induced Aβ-CRY2-mCh. Fewer cellular vacuoles observed. **(c)** HEK cells treated with lithium chloride shows homogenous expression of Aβ-CRY2-mCh, whereby blue light illumination did not result in significant clustering of Aβ-CRY2-mCh.

**Figure S8.**
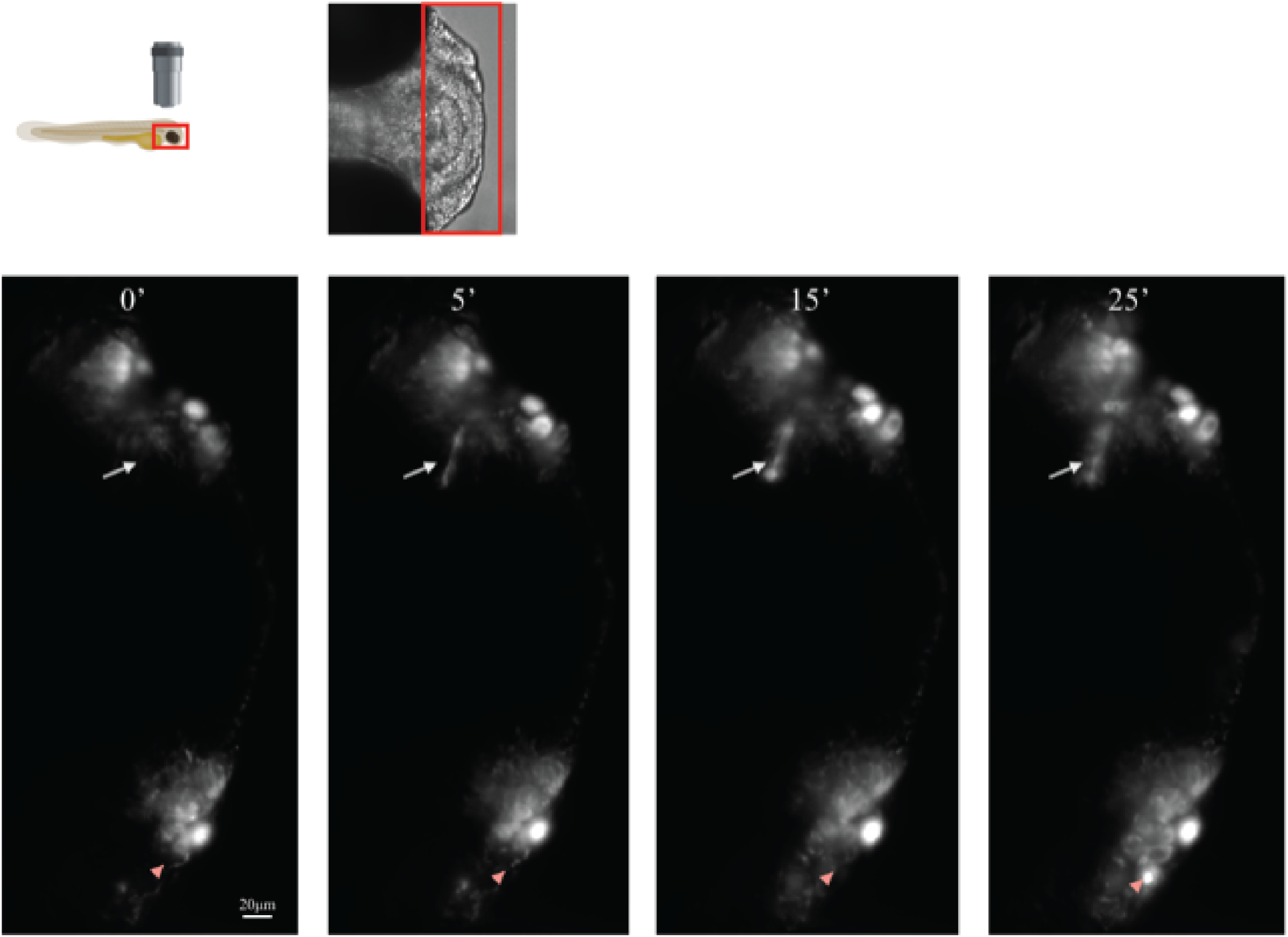
Transgenic Aβ model in *Danio Rerio embryo* expressing Aβ-CRY2-mCh (red).

## References

Ahn, N. G., T. S. Nahreini, N. S. Tolwinski and K. A. Resing (2001). “Pharmacologic inhibitors of MKK1 and MKK2.” Methods Enzymol 332: 417–431.

Anderson, R. M., C. Hadjichrysanthou, S. Evans and M. M. Wong (2017). “Why do so many clinical trials of therapies for Alzheimer’s disease fail?” The Lancet 390(10110): 2327–2329.

Antzutkin, O. N., D. Iuga, A. V. Filippov, R. T. Kelly, J. Becker-Baldus, S. P. Brown and R. Dupree (2012). “Hydrogen Bonding in Alzheimer’s Amyloid-β Fibrils Probed by 15N{17O} REAPDOR Solid-State NMR Spectroscopy.” Angewandte Chemie International Edition 51(41): 10289–10292.

Bischof, J., R. K. Maeda, M. Hediger and F. Karch (2007). “An optimized transgenesis system for Drosophila using germ-line-specific phiC31 integrases.” 104(9): 3312–3317.

Brand, A. H. and N. Perrimon (1993). “Targeted gene expression as a means of altering cell fates and generating dominant phenotypes.” Development 118(2): 401–415.

Butterfield, D. A., A. M. Swomley and R. Sultana (2013). “Amyloid beta-peptide (1-42)-induced oxidative stress in Alzheimer disease: importance in disease pathogenesis and progression.” Antioxid Redox Signal 19(8): 823–835.

Colosimo, P. F. and N. S. Tolwinski (2006). “Wnt, Hedgehog and junctional Armadillo/beta-catenin establish planar polarity in the Drosophila embryo.” PLoS One 1: e9.

Cummings, J. (2018). “Lessons Learned from Alzheimer Disease: Clinical Trials with Negative Outcomes.” Clin Transl Sci 11(2): 147–152.

Cummings, J., G. Lee, A. Ritter and K. Zhong (2018). “Alzheimer’s disease drug development pipeline: 2018.” Alzheimers Dement (N Y) 4: 195–214.

De Ferrari, G. V., M. A. Chacón, M. I. Barría, J. L. Garrido, J. A. Godoy, G. Olivares, A. E. Reyes, A. Alvarez, M. Bronfman and N. C. Inestrosa (2003). “Activation of Wnt signaling rescues neurodegeneration and behavioral impairments induced by β-amyloid fibrils.” Molecular Psychiatry 8: 195.

De-Paula, V. J., M. Radanovic, B. S. Diniz and O. V. Forlenza (2012). “Alzheimer’s disease.” Subcell Biochem 65: 329–352.

Duff, K., C. Eckman, C. Zehr, X. Yu, C. M. Prada, J. Perez-tur, M. Hutton, L. Buee, Y. Harigaya, D. Yager, D. Morgan, M. N. Gordon, L. Holcomb, L. Refolo, B. Zenk, J. Hardy and S. Younkin (1996). “Increased amyloid-beta42(43) in brains of mice expressing mutant presenilin 1.” Nature 383(6602): 710–713.

Fenno, L., O. Yizhar and K. Deisseroth (2011). “The development and application of optogenetics.” Annu Rev Neurosci 34: 389–412.

Ferreira, S. T. and W. L. Klein (2011). “The Abeta oligomer hypothesis for synapse failure and memory loss in Alzheimer’s disease.” Neurobiol Learn Mem 96(4): 529–543.

Fong, S., L. F. Ng, L. T. Ng, P. K. Moore, B. Halliwell and J. Gruber (2017). “Identification of a previously undetected metabolic defect in the Complex II Caenorhabditis elegans mev-1 mutant strain using respiratory control analysis.” Biogerontology 18(2): 189–200.

Fong, S., E. Teo, L. F. Ng, C.-B. Chen, L. N. Lakshmanan, S. Y. Tsoi, P. K. Moore, T. Inoue, B. Halliwell and J. Gruber (2016). “Energy crisis precedes global metabolic failure in a novel Caenorhabditis elegans Alzheimer Disease model.” Scientific Reports 6: 33781.

Fong, S., E. Teo, L. F. Ng, C. B. Chen, L. N. Lakshmanan, S. Y. Tsoi, P. K. Moore, T. Inoue, B. Halliwell and J. Gruber (2016). “Energy crisis precedes global metabolic failure in a novel Caenorhabditis elegans Alzheimer Disease model.” Sci Rep 6: 33781.

Forloni, G. and C. Balducci (2018). “Alzheimer’s Disease, Oligomers, and Inflammation.” J Alzheimers Dis 62(3): 1261–1276.

Groth, A. C., M. Fish, R. Nusse and M. P. Calos (2004). “Construction of Transgenic Drosophila by Using the Site-Specific Integrase From Phage ϕC31.” Genetics 166(4): 1775.

Guglielmi, G., J. D. Barry, W. Huber and S. De Renzis (2015). “An Optogenetic Method to Modulate Cell Contractility during Tissue Morphogenesis.” Dev Cell 35(5): 646–660.

Hardy, J. A. and G. A. Higgins (1992). “Alzheimer’s disease: the amyloid cascade hypothesis.” Science 256(5054): 184–185.

Hawkes, N. (2016). “Sixty seconds on … solanezumab.” BMJ 355.

Hong, E., K. Santhakumar, C. A. Akitake, S. J. Ahn, C. Thisse, B. Thisse, C. Wyart, J. M. Mangin and M. E. Halpern (2013). “Cholinergic left-right asymmetry in the habenulo-interpeduncular pathway.” Proc Natl Acad Sci U S A 110(52): 21171–21176.

Huang, A., C. Amourda, S. Zhang, N. S. Tolwinski and T. E. Saunders (2017). “Decoding temporal interpretation of the morphogen Bicoid in the early Drosophila embryo.” Elife 6.

Idevall-Hagren, O., E. J. Dickson, B. Hille, D. K. Toomre and P. De Camilli (2012). “Optogenetic control of phosphoinositide metabolism.” Proc Natl Acad Sci U S A 109(35): E2316–2323.

Iijima, K., H.-C. Chiang, S. A. Hearn, I. Hakker, A. Gatt, C. Shenton, L. Granger, A. Leung, K. Iijima-Ando and Y. Zhong (2008). “Aβ42 Mutants with Different Aggregation Profiles Induce Distinct Pathologies in Drosophila.” PLoS One 3(2): e1703.

Jin, N., H. Zhu, X. Liang, W. Huang, Q. Xie, P. Xiao, J. Ni and Q. Liu (2017). “Sodium selenate activated Wnt/beta-catenin signaling and repressed amyloid-beta formation in a triple transgenic mouse model of Alzheimer’s disease.” Exp Neurol 297: 36–49.

Johnson, H. E., Y. Goyal, N. L. Pannucci, T. Schupbach, S. Y. Shvartsman and J. E. Toettcher (2017). “The Spatiotemporal Limits of Developmental Erk Signaling.” Dev Cell 40(2): 185–192.

Kaur, P., T. E. Saunders and N. S. Tolwinski (2017). “Coupling optogenetics and light-sheet microscopy, a method to study Wnt signaling during embryogenesis.” Sci Rep 7(1): 16636.

Kennedy, M. J., R. M. Hughes, L. A. Peteya, J. W. Schwartz, M. D. Ehlers and C. L. Tucker (2010). “Rapid blue-light-mediated induction of protein interactions in living cells.” Nat Methods 7(12): 973–975.

Klein, P. S. and D. A. Melton (1996). “A molecular mechanism for the effect of lithium on development.” Proc Natl Acad Sci U S A 93(16): 8455–8459.

Kumar, A., A. Singh and Ekavali (2015). “A review on Alzheimer’s disease pathophysiology and its management: an update.” Pharmacol Rep 67(2): 195–203.

LaFerla, F. M., K. N. Green and S. Oddo (2007). “Intracellular amyloid-beta in Alzheimer’s disease.” Nat Rev Neurosci 8(7): 499–509.

Lim, H. C. and S. A. Mathuru (2018). “Modeling Alzheimer’s and Other Age Related Human Diseases in Embryonic Systems.” Journal of Developmental Biology 6(1).

Markstein, M., C. Pitsouli, C. Villalta, S. E. Celniker and N. Perrimon (2008). “Exploiting position effects and the gypsy retrovirus insulator to engineer precisely expressed transgenes.” Nature Genetics 40: 476.

Mroczko, B., M. Groblewska, A. Litman-Zawadzka, J. Kornhuber and P. Lewczuk (2018). “Amyloid beta oligomers (AbetaOs) in Alzheimer’s disease.” J Neural Transm (Vienna) 125(2): 177–191.

Müller, H. A. and E. Wieschaus (1996). “armadillo, bazooka, and stardust are critical for early stages in formation of the zonula adherens and maintenance of the polarized blastoderm epithelium in Drosophila.” The Journal of Cell Biology 134(1): 149.

Nyamsuren, O., D. Faggionato, W. Loch, E. Schulze and R. Baumeister (2007). “A mutation in CHN-1/CHIP suppresses muscle degeneration in Caenorhabditis elegans.” Dev Biol 312(1): 193–202.

Okazaki, A., Y. Sudo and S. Takagi (2012). “Optical silencing of C. elegans cells with arch proton pump.” PLoS One 7(5): e35370.

Park, J., I. Wetzel, I. Marriott, D. Dréau, C. D’Avanzo, D. Y. Kim, R. E. Tanzi and H. Cho (2018). “A 3D human triculture system modeling neurodegeneration and neuroinflammation in Alzheimer’s disease.” Nature Neuroscience 21(7): 941–951.

Parr, C., N. Mirzaei, M. Christian and M. Sastre (2015). “Activation of the Wnt/β-catenin pathway represses the transcription of the β-amyloid precursor protein cleaving enzyme (BACE1) via binding of T-cell factor-4 to BACE1 promoter.” The FASEB Journal 29(2): 623–635.

Rival, T., R. M. Page, D. S. Chandraratna, T. J. Sendall, E. Ryder, B. Liu, H. Lewis, T. Rosahl, R. Hider, L. M. Camargo, M. S. Shearman, D. C. Crowther and D. A. Lomas (2009). “Fenton chemistry and oxidative stress mediate the toxicity of the beta-amyloid peptide in a Drosophila model of Alzheimer’s disease.” The European journal of neuroscience 29(7): 1335–1347.

Stiernagle, T. “Maintenance of C. elegans.”

Toledo, E. M. and N. C. Inestrosa (2009). “Activation of Wnt signaling by lithium and rosiglitazone reduced spatial memory impairment and neurodegeneration in brains of an APPswe/PSEN1ΔE9 mouse model of Alzheimer&#39s disease.” Molecular Psychiatry 15: 272.

Tolwinski, N. S. and E. Wieschaus (2001). “Armadillo nuclear import is regulated by cytoplasmic anchor Axin and nuclear anchor dTCF/Pan.” Development 128(11): 2107–2117.

Tsujimoto, T., Y. Nakase and T. Sugano (1970). “Firefly luminescence for ATP assay.” Wakayama Med Rep 14(1): 29–32.

Westfall, S., N. Lomis and S. Prakash (2018). “Longevity extension in Drosophila through gut-brain communication.” Sci Rep 8(1): 8362.

Zhang, Y. W., R. Thompson, H. Zhang and H. Xu (2011). “APP processing in Alzheimer’s disease.” Mol Brain 4: 3.

